# Sufficiency of unidirectional allostery in KaiC in generating the cyanobacterial circadian rhythm

**DOI:** 10.1101/2020.04.01.021055

**Authors:** Shin-ichi Koda, Shinji Saito

## Abstract

The clock protein of cyanobacteria KaiC forms a homohexamer with two ring-shaped domains, C1 and C2. These domains undergo several domain-specific conformational transitions and allosterically communicate to generate a circadian rhythm. Interestingly, experiments show a possibility that C2 is independent of C1. However, detailed interplay among them remains elusive. Here we propose a mathematical model, which explicitly considers the interplay. The allostery in KaiC is here modeled to be unidirectional from C2 to C1. We demonstrate that the unidirectional allostery is sufficient for the circadian rhythm by showing the quantitative reproducibility of various experimental data, including temperature dependence of both phosphorylation oscillation and ATPase activity. Based on the present model, we further discuss possible functional roles of the unidirectional allostery particularly in the period robustness against both protein concentration and temperature.

## INTRODUCTION

Circadian clocks are biological timing systems embedded in most living organisms and enable the organisms to anticipate daily changes in the environment and to adjust their biological activities. The simplest circadian clock is that of cyanobacteria, where the core oscillator is composed of only three proteins, KaiA, KaiB, and KaiC[1]. Interestingly, this circadian rhythm can be reconstituted in a test tube just by mixing the three proteins with ATP, which results in a nearly 24-hour periodic oscillation of phosphorylation of KaiC[2]. In addition to this self-sustaining oscillation, the KaiABC oscillator possesses several fundamental functions as a biological clock. For example, the period of the oscillator is robust against environmental perturbations such as temperature[2] and concentrations of the proteins[3]. The oscillator is further entrained by various periodic environmental changes, including temperature[4] and ATP/ADP ratio in buffer[5]. This simple yet functional system has thus attracted considerable interests in elucidating the molecular origins of circadian rhythm.

KaiC is the central component in the KaiABC oscillator, and in the presence of ATP, forms a homohexamer with two ring-shaped domains, C1 and C2[6, 7]. Both C1 and C2 have ATPase activities. C2 also has autokinase and autophosphatase activities, where two residues near the ATP binding site in C2, Ser431 and Thr432, are phosphorylated and dephosphorylated. With the help of KaiA and KaiB, (de)phosphorylation occurs in the following order: ST → SpT → pSpT → pST → ST[8, 9], where S/pS and T/pT are the unphosphorylated/phosphorylated states of Ser431 and Thr432, respectively.

KaiA promotes phosphorylation of KaiC[10, 11] by acting on the C-terminal tail of C2[12–15]. It facilitates the exchange of bound ADP with exogenous ATP[16], suggesting that the enhanced ADP/ATP exchange supplies abundant phosphate for phosphorylation reactions in the form of ATP, and shifts the equilibrium of phosphorylation reactions toward phosphorylated states.

KaiB, on the other hand, switches the oscillation phase to dephosphorylation by inhibiting KaiA activity[17, 18]. When C2 is adequately phosphorylated, KaiB binds to C1 and strongly sequesters KaiA from C2 by forming C1 *·* KaiB *·* KaiA complex[19]. Indeed, this complex has recently been observed experimentally[20, 21]. After this sequestration, KaiA no longer promotes phosphorylation, and C2 starts to dephosphorylate, thus the phosphorylation oscillation occurs.

During the oscillation, C1 and C2 allosterically communicate and regulate the (dis)assembly of KaiA and KaiB through several domain-specific conformational transitions. For example, recent studies on the KaiABC complex[20, 21] have revealed that KaiB can bind to an ADP-bound conformation of C1[22]. On the other hand, C2 has the buried and exposed states of the C-terminal tail[14, 15], and KaiA can bind only in the latter state. C2 also takes the rigid and flexible conformational states that regulate C1-C2 stacking[19, 23]. Yet, it remains elusive how these multiple conformational transitions interplay each other and link the chemical reactions of ligands, e.g. ATP hydrolysis and (de)phosphorylation reaction, with the (dis)assembly of Kai proteins. In particular, we here focus on an unaccounted experimental result that modulations on C1 do not show apparent effect on the dephosphorylation process in C2[24, 25]. Interestingly, this C1-independent C2 dephosphorylation contradicts an intuition that C1 and C2 should bidirectionally interact to generate the oscillation.

In this article, we conduct a quantitative investigation with a novel mathematical reaction model to clarify the relation between the functions of the KaiABC oscillator and the interplay among the domain-specific conformational transitions. Our model has four main features below. First, the model explicitly considers multiple domain-specific conformational transitions and the interplay among them. In particular, we hypothetically design the allosteric communication in KaiC to be unidirectional from C2 to C1 so that the model can realize the C1-independent C2 phosphorylation mentioned above. In contrast, previous models considering conformational states[26–29] have assumed only a single global conformational transition of overall KaiC. This single global conformational transition, where C1 and C2 bidirectionally interplay, could be an oversimplified assumption because further assumptions are needed to realize the C1-independent C2 dephosphorylation. Second, in contrast to conceptually simplified models, the present model is described as a set of elementary reactions, which consists of the chemical reactions of ligands, (dis)assembly of KaiA and KaiB, and the conformational transitions of KaiC. With this representation, the present model allows detailed investigations of the clock functions on molecular basis. Third, owing to the representation based on elementary reactions, an Arrhenius temperature dependence is introduced to rate or equilibrium constant of each elementary reaction, which enables investigations on the temperature compensation of the period. Lastly, the parameter values of the present model are determined by an automatic optimization, which encourages the reproducibility of several fundamental experimental data. This approach reduces arbitrariness in the parameter values compared to the manually determined values, which are often used to explain a specific experimental result. Thus, the present parameter values improve the reliability of the model. To our knowledge, there has been no previous model satisfying all the features above.

With the present model, we investigate the relation between the elementary reactions and clock functions of the KaiABC oscillator. In particular, we focus on the hypothetical assumption of the unidirectional allostery from C2 to C1. We first show that the unidirectional allostery is sufficient to generate the circadian rhythm by demonstrating that the present model can indeed reproduce various experimental results, including temperature dependence of both phosphorylation oscillation and ATPase activity. Based on the present model, we further discuss possible functional roles of the unidirectional allostery in the generation of collective oscillation and in the period robustness against both protein concentration and temperature.

## RESULTS

### Model setup

The present model consists of elementary reactions described in this subsection, which is organized as follows. We first describe the processes at each ATP binding site of C1 and C2. Then, we summarize the domain-specific conformational transitions of C1 and C2, together with the (dis)assembly of KaiA and KaiB. Lastly, we explain the allosteric communication between C1 and C2 in the present model, which is unidirectional from C2 to C1. The details of the model are found in Method.

At each C1 ATP binding site in the present model, only ATP hydrolysis and the ADP/ATP exchange are considered. ATP synthesis is neglected as it is supposed to be very slow due to a high energy barrier. The effect of inorganic phosphate produced by ATP hydrolysis is also ignored for simplicity. The ADP/ATP exchange actually consists of two successive processes, i.e. the release of bound ADP and incorporation of exogenous ATP. In the present model, however, these two processes are combined and referred to as the ADP/ATP exchange because the incorporation of ATP is rapid[7].

The processes at each C2 ATP binding site in the present model are summarized in Fig. 1. We use the following notations for phosphorylation and bound-nucleotide states: U (unphosphorylated); S or T (either Ser431 or Thr432 is phosphorylated, respectively); and D (doubly phosphorylated), with a subscript T or D to indicate a bound nucleotide, ATP or ADP, respectively. In C2, we consider ATP hydrolysis and ADP/ATP exchange as in C1. Moreover, we consider phosphorylation and dephosphorylation reactions of Ser431 and Thr432, in which a phosphate is transferred between the bound nucleotide and either of Ser431 and Thr432[30, 31]. Note that the processes in each ATP binding site in KaiC are the same as Paijmans’s model[29].

**FIG. 1.**
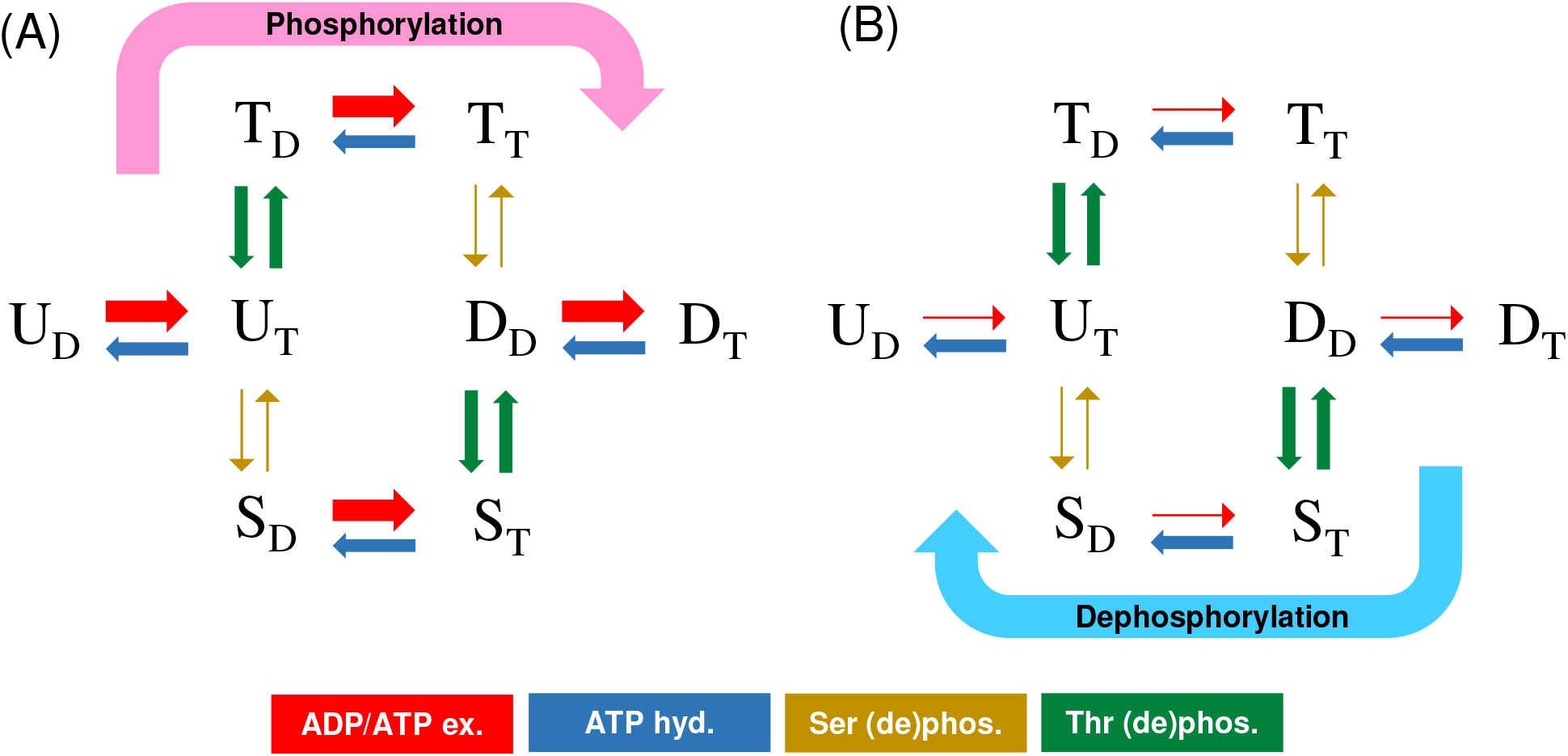
Reaction scheme at a C2 ATP binding site. Red and blue arrows represent the ADP/ATP exchange and ATP hydrolysis (with phosphate release), and ocher and green arrows represent (de)phosphorylation of Ser431 and Thr432, respectively. Arrow thickness qualitatively represents the magnitude of a rate constant. The ADP/ATP exchange is the largest in (A) and the smallest in (B).

It has been experimentally suggested that the change of the ADP/ATP exchange in C2 induces an equilibrium shift toward (de)phosphorylated states of C2[16]. The present model designs the sequential phosphorylation of C2 as schematically shown in Fig. 1, in which the equilibrium shift is drawn by thick arrows. Assuming that (de)phosphorylation of Thr432 is faster than that of Ser431, an enhanced ADP/ATP exchange promotes the phosphorylation via the T state (Fig. 1A), whereas a suppressed ADP/ATP exchange results in the dephosphorylation via the S state (Fig. 1B).

In the present model, one C1-specific conformational transition, between KaiB-binding-competent (BC) and -incompetent (BI) conformational states, is considered (Fig. 2A). We assume that KaiB can bind to only C1 in the BC state and that KaiA can bind strongly to the KaiB-C1 complex, which eventually results in the sequestration of KaiA from C2. Moreover, the conformational transition from BI to BC state is assumed to be promoted when C1 binds abundant ADP rather than ATP, as suggested experimentally[20, 21, 32, 33]. Under this assumption, switching between phosphorylation and dephosphorylation is controlled by the ADP/ATP exchange in C1 because the exchange determines the amount of ADP in C1, which is the trigger of the conformational transition from BI to BC state. Specifically, a suppressed ADP/ATP exchange accumulates ADP molecules from ATP hydrolysis and consequently increases the BC state of C1 and the KaiA-KaiB-C1 complex. This mechanism is consistent with the recent experimental result that the KaiB binding requires the ATP hydrolysis of C1 in advance[34].

**FIG. 2.**
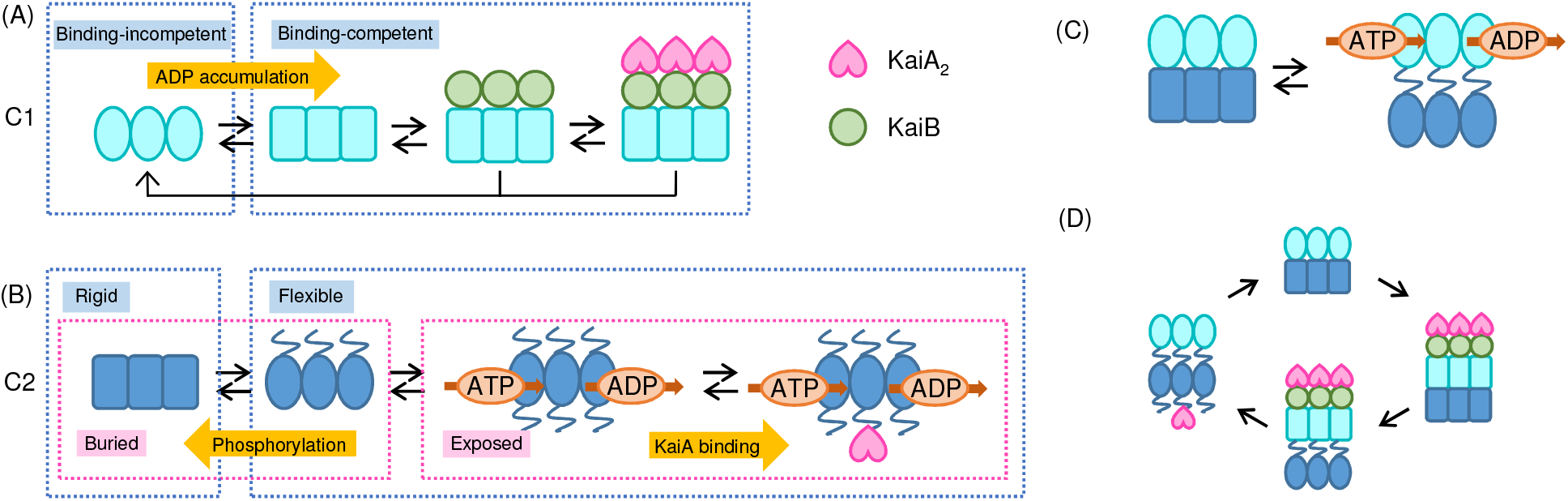
Conformational transitions of KaiC in the present model. (A) Conformational transition of C1. Schematic figures of C1 with elliptical and rectangular shapes represent KaiB-binding-incompetent (BI) and -competent (BC) conformational states, respectively. KaiB can bind to C1 in the BC state, and KaiA can bind to KaiB bound to C1. (B) Conformational transitions of C2. Schematic figures of C2 with rectangular and elliptical shapes represent rigid and flexible conformational states of C2, respectively. Strings drawn from the top of C2 (linkers between C1 and C2) correspond to C1-C2 stacking, which is assumed to occur along with C2 rigidification. Strings drawn from the bottom of C2 (C-terminal tails) represent buried and exposed conformational states of C2. The exposed state, which is absent in the rigid state, can bind KaiA and allows ADP/ATP exchange in C2. (C) Allosteric communication in KaiC. C2 in the rigid state prohibits the ADP/ATP exchange in C1 due to the rigidification of C2. (D) Cycle of conformational transitions of KaiC during phosphorylation oscillation.

In addition, two types of C2-specific conformational transitions are considered (Fig. 2B): one is between the rigid and flexible conformational states of C2[23], and the other is between the buried and exposed conformational states of the C-terminal tail[14, 15]. Based on the two experimental results showing that the phosphorylation of C2 rigidifies C2[23] and buries the tail[15], we assume that the exposed state of the tail is absent when C2 is in the rigid state. Hence, only three combinational states, i.e. rigid-buried, flexible-buried, and flexible-exposed states, are considered. Moreover, we assume that KaiA can bind to C2 only when C2 is in the exposed state. To realize the experimental result that KaiA facilitates the ADP/ATP exchange in C2[16], we also assume that the ADP/ATP exchange in C2 is allowed only in the exposed state. With these assumptions, the KaiA binding promotes the ADP/ATP exchange through the stabilization of the exposed state, i.e. the equilibrium shift of Fig. 2B to right.

For the allosteric communication between C1 and C2 in the present model, we hypothetically assume that C2 in the rigid conformational state prohibits the ADP/ATP exchange in C1. This idea originates from our assumption that the C1-C2 stacking, which concurrently occurs with the rigidification of C2[19, 23], prohibits the ADP/ATP exchange. In the present model, however, we do not explicitly describe the C1-C2 stacking for simplicity. We here emphasize that this allostery is unidirectional, i.e. the status of C1 does not affect C2. Thus, the C1-independent C2 dephosphorylation[19, 23] is realized without further assumptions in the present model.

Figure 2D illustrates the overview of the entire oscillation process in the present model explained above. Starting from KaiA-C2 binding (left figure of Fig. 2D), which stabilizes the exposed conformational state of C2 and promotes the ADP/ATP exchange and phosphorylation of C2, phosphorylated C2 rigidifies itself and induces C1-C2 stacking (upper figure). The rigid C2, in turn, inhibits the ADP/ATP exchange in C1 while facilitating ADP accumulation. ADP-rich C1 is then transformed from BI to BC conformational state of C1, which induces the formation of the C1-KaiB-KaiA complex, i.e. the sequestration of KaiA from C2 (right figure). As KaiA no longer acts on C2 due to the sequestration, C2 dephosphorylation proceeds along with the conformational transition to the flexible conformational state of C2 (lower figure), which allows the ADP/ATP exchange in C1. Finally, after the release of ADP from C1, C1 returns to the BI conformational state and releases KaiB and KaiA (return to the left figure again).

Here, we make some remarks on KaiA and KaiB. All KaiA proteins are treated as dimers in this model. Although KaiB proteins are known to form monomers, dimers, and tetramers, all KaiB proteins are treated as monomers in this model for simplicity. Indeed, monomeric form is dominant near the so-called standard condition (3.5 *μ*M)[35]. A recent experiment has shown that KaiB has two conformational states, and the transition between them plays a role in the slow KaiB-C1 binding[36]. In the present model, however, we consider only one conformational state of KaiB to focus more on the conformational transitions of KaiC and assume that the slowness arises from C1 rather than KaiB, as claimed by a recent experimental study[34].

### Parameter Optimization

Next, as a validation of the present model, we show that the present model can quantitatively reproduce various experimental data. The parameter values are determined by an automatic optimization method that simultaneously fits multiple types of model outputs to corresponding experimental data. Here, we optimize 19 independent rate and equilibrium constants with an Arrhenius temperature dependence (38 parameters, i.e. their pre-exponential factor, the activation energy of rate constants, and the activation energy difference of equilibrium constants between corresponding forward and backward rate constants). The detailed procedure of the optimization, the experimental data used in the optimization, and the optimized parameter values are summarized in Method.

Figure 3 shows the results of the fitting. Figures 3A-D exhibit the phosphorylation oscillation of KaiC at 26-38 °C. In these cases, the results of the model are in good agreement with the corresponding experimental data[37], which indicates that the temperature compensation of the period is realized in the present model. Figures 3E and F show the phosphorylation of KaiC in the absence of both KaiA and KaiB (Fig. 3E) and in the presence of KaiA (Fig. 3F), in which the experimental data of the U and T states in the former case[38] at time *t* = 20 h and the data of the four states in the latter case[8] at time *t* = 20 h are used in the optimization. In these cases, the results of the model slightly deviate from the experimental data used in the optimization. However, relaxation behaviors, e.g. over-shoots of the S state in the absence of KaiAB (Fig. 3E) and of the T state in the presence of KaiA (Fig. 3E), are well reproduced although the experimental data are adopted only from a time point (*t* = 20 h). Figure 3G shows the ratio of the bound ATP in KaiC to the bound nucleotide, in which the corresponding experimental data[16] at time *t* = 8 h are considered in the optimization. Although the result of the model in the presence of KaiA deviates from the experimental data to some extent, the model reproduces the experimental result that the ratio in the presence of KaiA is larger than in the absence of KaiAB due to the promotion of the ADP/ATP exchange in C2 by KaiA[16]. Figure 3H shows the ATPase activities, i.e. the rates of ADP production, in the absence of KaiAB at 25-40 °C. The result of the model exhibits the initial temperature dependence (thermal sensitivity Q_10_ = 1.3 at *t* = 0.1 h) and the temperature compensation at the steady state (Q_10_ = 1.1 at *t* = *∞*), which are consistent with the experimental results[22] used in the optimization (Q_10_ = 1.4 and 1.1, respectively). Moreover, the present model can reproduce the overshoot of the ATPase activity[22] without additional experimental data. Figure 3I shows the total KaiA concentration dependence of the period in the present model, in which the experimental data[3] at the total concentration A_tot_ = 3.6 *μ*M is used in the optimization. The period of the model is almost constant when A_tot_ *<* 3.5 *μ*M as in the experiments. The total KaiB concentration dependence of the phosphorylation oscillation[3] is also well reproduced by the present model (Fig. 3J); the phosphorylation oscillation is almost independent of the total concentration B_tot_ when B_tot_ *>* 1.8 *μ*M, while the rhythmic behavior drastically disappears with decreasing in B_tot_ below 1.8 *μ*M. Other results of the fitting are shown in Supplementary Information.

**FIG. 3.**
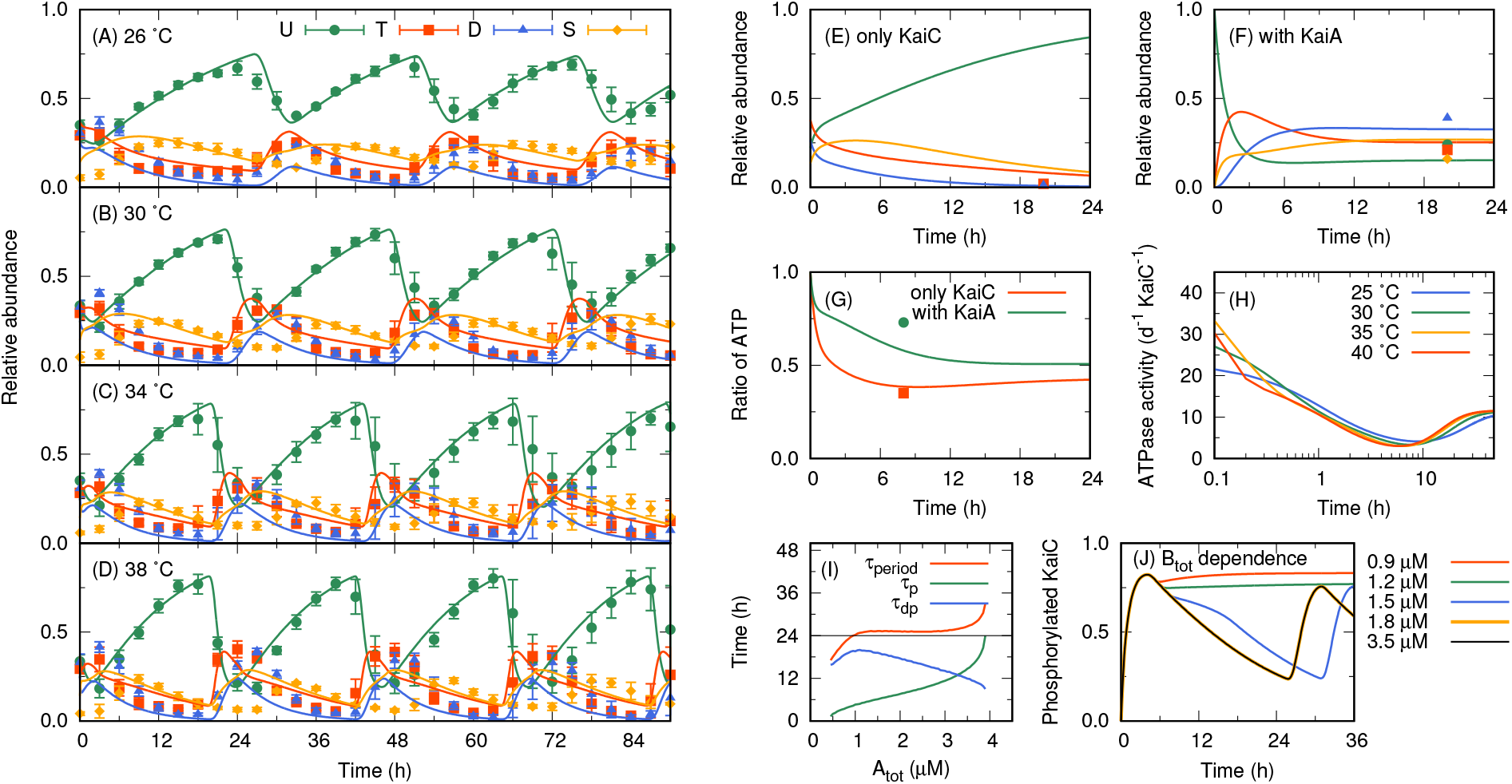
Results of the present model with optimized parameters. Solid curves and dots represent the results of the model and the experimental data used in the parameter optimization. (A-F) Relative abundance of phosphorylation states (U, T, D, and S) of KaiC. (A-D) show the relative abundance of the system in the presence of KaiAB at 26, 30, 34, and 38 °C. Dots and error bars indicate the experimental data and their standard deviations[37], respectively. The initial state of the model is the steady state in the absence of KaiAB at 0 °C. (E) shows the relative abundance of the system in the absence of KaiAB at 30 °C with the same initial state as that used in (A-D). The experimental data are taken from Ref. [38]. (F) shows the relative abundance of the system in the presence of KaiA at 30 °C. The initial state is fully dephosphorylated and binds ATP. The experimental data are taken from Ref. [8]. (G) Ratios of bound ATP in KaiC to bound nucleotide at 30 °C in the absence of KaiAB and in the presence of KaiA. The initial state is the same as that used in (F). The experimental data are taken from Ref. [16]. (H) Temperature dependence of ATPase activity in the absence of KaiAB. The initial phosphorylation state of the model is the steady state in the absence of KaiAB at 0 °C, but all bond nucleotides are replaced to ATP. The experimental result of thermal sensitivities *Q*_10_ at time *t* = 0.1 h and *∞*[22] are used in the optimization (not shown). (I) A_tot_ dependences of the period *τ*_period_, the duration of phosphorylation *τ*_p_, and that of dephosphorylation *τ*_dp_ at 30 °C. (J) B_tot_ dependence of the phosphorylation oscillation. The initial state is the same as that used in (F). The experimental data from Ref. [3] are used in the optimization to obtain (F) and (J) (Method).

### Intra-KaiC unidirectional allostery

Next, we investigate the origins of the clock functions of the KaiABC oscillator with the present model. In particular, we here focus on the unidirectional allostery from C2 to C1 in the present model, which is hypothetically introduced to realize the C1-independent C2 dephosphorylation[24, 25], and discuss the functional roles of the allostery on the generation of oscillation and on the period robustness against environmental perturbations.

Although the unidirectional allostery apparently contradicts with an intuition that C1 and C2 may bidirectionally interplay, the present model indeed shows that the unidirectional allostery is sufficient to generate a circadian rhythm. This sufficiency is assured by KaiA surrounding KaiC because C1 can indirectly communicate with C2 through the concentration of unbound KaiA, which is controlled by the sequestration of KaiA at C1 (Fig. 4); in the present model, information on phosphorylation is unidirectionally transmitted from C2 to C1 by the allostery of each KaiC hexamer, whereas information on phosphorylation/dephosphorylation switching is indirectly transmitted from C1 to C2 via the concentration of the free KaiA surrounding KaiC.

**FIG. 4.**
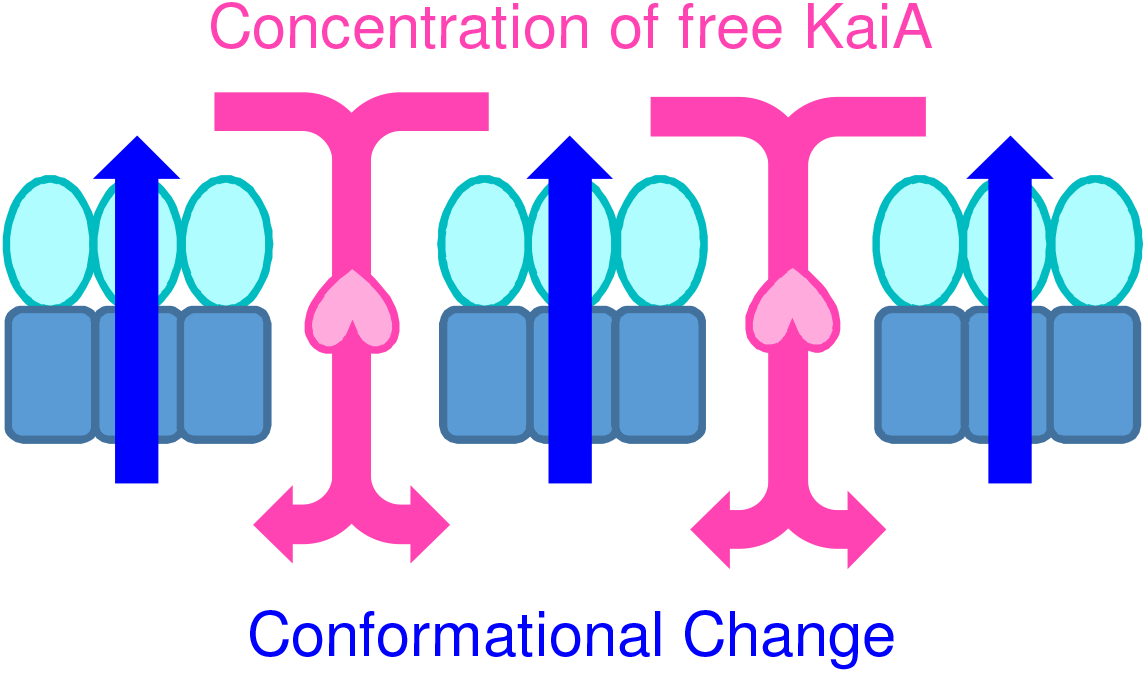
Direction of information transmission in the present model. Information on the phosphorylation state is transmitted from C2 to C1 by conformational transitions of each KaiC hexamer (blue arrows), whereas information on phosphorylation/dephosphorylation switching is indirectly transmitted from C1 to C2 by the concentration of free KaiA surrounding KaiC (red arrows).

As discussed previously[26, 29], the “collectivity” of the KaiABC oscillator may be given by the “inter”-KaiC communication mediated by KaiA: phosphorylated KaiC sequesters KaiA and waits for other unphosphorylated KaiC to be phosphorylated; after free KaiA is decreased below a threshold due to the sequestration, dephosphorylation of KaiC occurs at once. We here make a remark that the unidirectionality of the allostery in each KaiC hexamer may have a positive role on an effective generation of the collective oscillation. If C1 and C2 bidirectionally interplay, the bidirectional communication could switch the oscillation phase within a hexamer independently of other hexamers, and hence, the waiting process mentioned above could be disrupted.

### Period robustness against the total concentration of KaiA

As shown above, the present model can reproduce the period robustness against the total concentration of KaiA (Fig. 3I) and temperature (Fig. 3A-D). To explain possible mechanisms of the robustness in terms of the allosteric communication between C1 and C2, we here focus on the attenuation of C1-KaiB binding by KaiA acting on C2. This attenuation has been observed experimentally and was originally modeled through a global conformational transition of the whole KaiC[28], where C1 and C2 bidirectionally communicate. However, it is noted that this attenuation is also realized in the present model with only the unidirectional allostery. Specifically, KaiA binding to C2 stabilizes the flexible conformational state of C2 by shifting the equilibrium shown in Fig. 2B to right. This flexible state then triggers the ADP/ATP exchange in C1, which prevents the KaiB binding to C1 as explained in the model setup.

To investigate the origin of the period robustness against the total concentration of KaiA, A_tot_, we first divide the period of phosphorylation oscillation into the durations of phosphorylation *τ*_p_ and dephosphorylation *τ*_dp_. Here *τ*_p_ and dephosphorylation *τ*_dp_ are defined by the period when more than 0.5 percent of KaiA binds to C2 or not, respectively. In the present model, *τ*_dp_ decreases, whereas *τ*_p_ increases with increasing A_tot_ (Fig.3I).

The decrease of *τ*_dp_ with increasing A_tot_ can be explained as follows. As suggested in many previous studies, KaiA is sufficiently sequestered from C2 by KaiB bound to C1 in the dephosphorylation phase. The oscillation then switches to the phosphorylation phase when the amount of KaiB bound to C1 falls below the threshold for the sufficient KaiA sequestration. Assuming this mechanism, the increase in A_tot_ lifts the threshold because KaiB must sequester more KaiA, and hence, the dephosphorylation phase terminates earlier.

On the other hand, the increase in *τ*_p_ with increasing A_tot_ can be attributed to the attenuation of C1-KaiB binding by KaiA acting on C2 mentioned above. With increasing A_tot_, the attenuated KaiB binding to C1 delays the subsequent KaiA sequestration, and hence *τ*_p_ becomes long.

The period robustness against KaiA concentration can be explained by a proper balance between the opposing effects on *τ*_p_ and *τ*_dp_. Indeed, with a modified parameter to disrupt the balance, the present model loses the period robustness (Figs. 5A and B). Specifically, an increase in the rate of ADP/ATP exchange of C1, *k*_1ex_, enhances the attenuation of KaiB binding by KaiA because the exchange inhibits the KaiB binding as explained in the model setup, and hence, *τ*_p_ becomes more sensitive to A_tot_ change with increasing *k*_1ex_ (Fig. 5B), and vice versa (Fig. 5A). The present model thus indicates that the proper balance is essential for the period robustness against A_tot_.

**FIG. 5.**
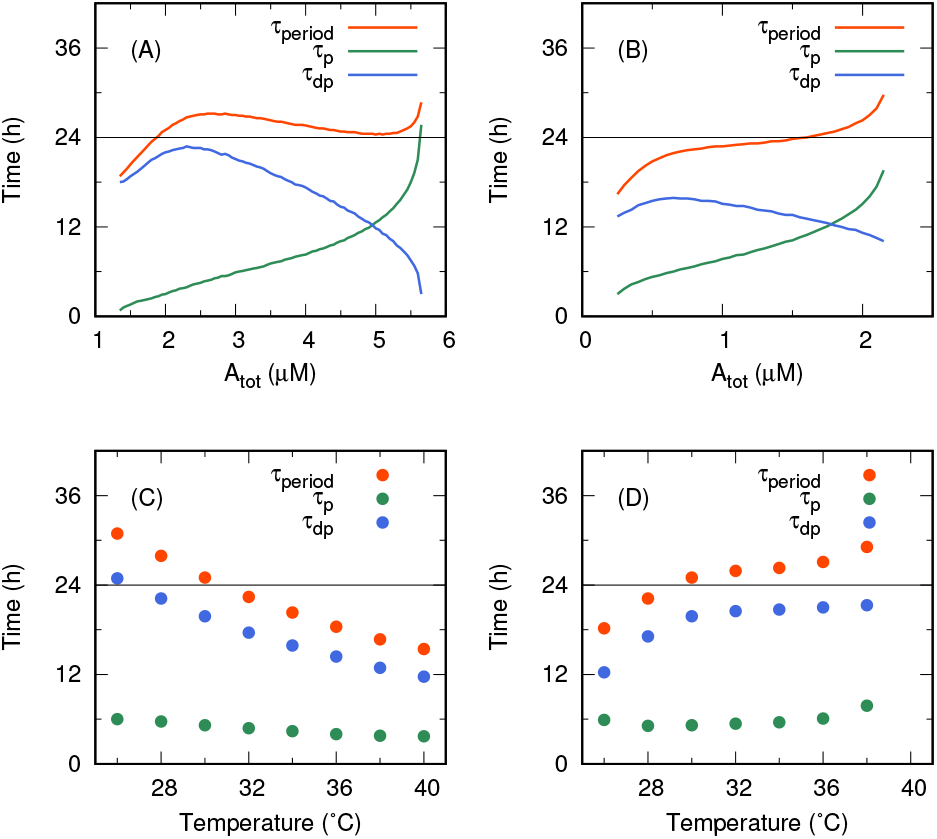
Environmental parameter dependences of the periods in the present model. A_tot_ dependences of *τ*_period_, *τ*_p_, and *τ*_dp_ at 30 °C (A) when *k*_1ex_ = 5 h^*−*1^ and (B) *k*_1ex_ = 30 h^*−*1^. Temperature dependences of *τ*_period_, *τ*_p_, and *τ*_dp_ (C) when the activation energy difference of *K*_2A_ is 0 kcal/mol and (D) is *−*80 kcal/mol. The pre-exponential factor of *K*_2A_ is determined so that *K*_2A_ coincides with its optimized value at 30 °C.

### Period robustness against temperature

Lastly, we discuss how the period robustness against temperature, i.e. the temperature compensation of the period, arises from the combination of elementary reactions. As experiments have shown that not only the period but also both *τ*_p_ and *τ*_dp_ are temperature-compensated (Figs. 3A-D), we separately discuss *τ*_p_ and *τ*_dp_ again.

The temperature compensation of *τ*_p_ in the present model can be explained by the balance between the following two opposing effects. On the one hand, *τ*_p_ is shortened with increasing temperature because the phosphorylation reactions are accelerated due to their high activation energies (see Table I. For example, 30.0 and 36.5 kcal/mol for the phosphorylation reactions of Ser431 and Thr432 in the rigid C2, respectively). On the other hand, *τ*_p_ is prolonged with increasing temperature because the attenuation of KaiB binding to C1 by KaiA is enhanced. The optimized parameters indeed show that the affinity between C2 and KaiA is strengthened when temperature increases; the equilibrium constant of C2-KaiA binding, *K*_2A_, decreases because its activation energy difference, i.e. coefficient of the temperature-dependent term in the exponent, is largely negative (*−*32.4 kcal/mol).

**TABLE I.** Optimized parameters

| Parameter | Value at 30 °C | Activation energy or<br>activation energy difference<br>(kcal/mol) | Constraint |
| --- | --- | --- | --- |
| $[A_2]_{\text{tot}}$ | $6.00 \times 10^{-1} \mu\text{M}$ | 0 | Fixed |
| $[B]_{\text{tot}}$ | $3.50 \times 10^0 \mu\text{M}$ | 0 | Fixed |
| $[C_6]_{\text{tot}}$ | $5.83 \times 10^{-1} \mu\text{M}$ | 0 | Fixed |
| $k_{1h}$ | $5.47 \times 10^{-1} \text{ h}^{-1}$ | $2.36 \times 10^0$ | |
| $k_{1ex}^{\text{BI F}}$ | $1.28 \times 10^1 \text{ h}^{-1}$ | $7.68 \times 10^{-1}$ | |
| $k_{1ex}^{\text{other}}$ | $0 \text{ h}^{-1}$ | 0 | Fixed |
| $k_{1IC,0}$ | $5.92 \times 10^{-3} \text{ h}^{-1}$ | $5.37 \times 10^{-1}$ | |
| $\delta\mu_{1ADP}$ | $1.41 \times 10^0$ | $2.95 \times 10^{-2}$ | |
| $k_{1CI}$ | $3.44 \times 10^0 \text{ h}^{-1}$ | $2.81 \times 10^0$ | |
| $K_{1B}$ | $1.0 \times 10^{-5} \mu\text{M}$ | 0 | Fixed |
| $K_{1A}$ | $1.0 \times 10^{-5} \mu\text{M}$ | 0 | Fixed |
| $k_{2h}$ | $2.49 \times 10^{-1} \text{ h}^{-1}$ | $7.07 \times 10^{-1}$ | |
| $k_{2ex}^B$ | $0 \text{ h}^{-1}$ | 0 | Fixed |
| $k_{2ex}^E$ | $6.77 \times 10^0 \text{ h}^{-1}$ | $9.86 \times 10^{-1}$ | |
| $k_{S+}^R$ | $1.74 \times 10^{-1} \text{ h}^{-1}$ | $3.00 \times 10^1$ | |
| $k_{S-}^R$ | $1.69 \times 10^{-1} \text{ h}^{-1}$ | $3.71 \times 10^1$ | |
| $k_{S+}^F$ | $5.73 \times 10^{-1} \text{ h}^{-1}$ | $5.05 \times 10^{-1}$ | |
| $k_{S-}^F$ | $1.31 \times 10^0 \text{ h}^{-1}$ | $6.27 \times 10^0$ | Constrained by Eq. (19) |
| $k_{T+}^R$ | $1.16 \times 10^0 \text{ h}^{-1}$ | $3.65 \times 10^1$ | |
| $k_{T-}^R$ | $1.32 \times 10^{-1} \text{ h}^{-1}$ | $4.04 \times 10^1$ | |
| $k_{T+}^F$ | $2.10 \times 10^{-1} \text{ h}^{-1}$ | $5.11 \times 10^0$ | |
| $k_{T-}^F$ | $1.76 \times 10^{-1} \text{ h}^{-1}$ | $1.72 \times 10^1$ | Constrained by Eq. (20) |
| $K_{2FR,0}^B$ | $9.12 \times 10^0$ | $-5.67 \times 10^0$ | |
| $\delta\mu_{2pS}$ | $1.08 \times 10^0$ | $-4.37 \times 10^0$ | |
| $\delta\mu_{2pT}$ | $-7.92 \times 10^{-2}$ | $5.16 \times 10^0$ | |
| $\delta\mu_{2ADP}$ | $-2.27 \times 10^{-1}$ | $3.02 \times 10^0$ | |
| $K_{2EB}^R$ | ∞ | 0 | Fixed |
| $K_{2EB}^F$ | $5.60 \times 10^2$ | $1.47 \times 10^1$ | |
| $K_{2A}$ | $1.14 \times 10^{-4} \mu\text{M}$ | $-3.24 \times 10^1$ | |

In the present model, an imbalance between the two opposing effects on *τ*_p_ indeed leads to a loss of the temperature compensation of *τ*_p_ (Figs. 5C and D). We here modify the temperature dependence of *K*_2A_ to disrupt the balance. When *K*_2A_ is assumed to be independent of temperature (Fig. 5C), the increase in *τ*_p_ is lost. In this case, *τ*_p_ decreases as temperature rises (*Q*_10_ = 0.67). On the other hand, when we amplify the temperature dependence of *K*_2A_ with a higher activation energy difference (*−*80 kcal/mol) (Fig. 5D), *τ*_p_ increases with temperature (*Q*_10_ = 1.53). Thus, in the present model, the temperature compensation of *τ*_p_ is described by the balance between the acceleration of phosphorylation reactions and the attenuation of C1-KaiB binding by KaiA.

The temperature compensation of *τ*_dp_, on the other hand, can be explained through the ratio of the rate constants of phosphorylation to dephosphorylation reactions. Because the amplitude of the phosphorylation oscillation can be approximated by 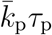 as well as by 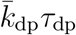, where 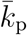 and 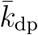 are the mean rate constants of the phosphorylation and dephosphorylation reactions, respectively, we obtain

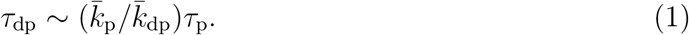

As *τ*_p_ is temperature-compensated, the temperature compensation of *τ*_dp_ is achieved when 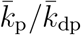 is also temperature-compensated, i.e. the activation energies of 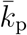 and 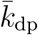 are in the same order. This is the case in the present model; for instance, the activation energies of phosphorylation reaction of Thr432 and dephosphorylation reaction of Ser431 in the rigid conformational state are 36.5 and 37.1 kcal/mol, respectively. Note that the validity of the estimation in Eq. (1) can be alternatively confirmed by the results shown in Figs. 5C and D. With Eq. (1) and the temperature competition of 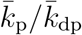, *τ*_p_*/τ*_dp_ is expected to be temperature-compensated even when each of *τ*_p_ and *τ*_dp_ depends on temperature. Indeed, *τ*_p_*/τ*_dp_ in Figs. 5C and D are relatively temperature-compensated (*Q*_10_ = 1.1 and 1.2, respectively).

## DISCUSSION

In this study, we developed a mathematical model of the circadian rhythm of Kai proteins consisting of the elementary reactions: chemical reactions of ligands, domain-specific conformational transitions, and (de)formation of complex among KaiABC. Most of the conventional mathematical models of the KaiABC oscillator have been conceptually simplified, i.e. represented by simplified processes rather than elementary reactions, to find universality in biological clocks. In contrast, the present model elucidate the microscopic origin of biological functions in the system by decomposing the behavior of the system into the elementary reactions.

The allosteric communication between C1 and C2 is crucial to generate the circadian rhythm. To investigate this allostery, we introduced the domain-specific conformational transitions in the present model, in contrast to a single global conformational transition of the whole KaiC assumed in the previous studies[26–29]. Importantly, we hypothetically designed the C2-dependent ADP/ATP exchange in C1 as the only explicit allosteric communication between C1 and C2. This allostery is unidirectional from C2 to C1, and thus explains the C1-independent dephosphorylation of C2[24, 25] without further assumptions. The present model reveals that the unidirectional allostery is sufficient for the generation of the oscillation and is important for the collectivity of the oscillation.

We further showed that the unidirectional allostery is sufficient to achieve the attenuation of C1-KaiB binding by the KaiA acting on C2[28]. In the present model, this attenuation yields the period robustness against both total KaiA concentration and temperature by adjusting the duration of the phosphorylation phase *τ*_p_. This result means that the period robustness against the two very different factors may arise from a common molecular origin via the unidirectional allostery.

There have been several studies on the period robustness against the total KaiA concentration[28, 39], which have suggested that the robustness requires the multimerization of KaiC[28] and the phosphorylation-dependent affinity between KaiA and C2[39]. In this respect, the present model showed that these factors are insufficient to achieve the period robustness; the present model, which considers both multimerization and phosphorylation-dependent affinity, can still lose the period robustness by disrupting the *τ*_p_ adjustment. Our result suggests that a properly balanced *τ*_p_ adjustment is also needed for the period robustness.

In the present study, we did not discuss the mechanism of entrainment to environmental changes, which is another fundamental property of biological clocks. This is to be discussed elsewhere.

## METHODS

### Multimeric forms of KaiABC in the present model

KaiC in the present model takes a hexameric form. Both C1 and C2 domains have six ATP binding sites and undergo their own conformational transitions. The total concentration of KaiC hexamer [C_6_]_tot_ is fixed to the so-called standard condition (3.5/6 *μ*M). KaiA and KaiB in the present model take dimeric and monomeric forms, respectively. Both KaiA and KaiB are assumed to have only one conformation. The total concentrations of KaiA dimer [A_2_]_tot_ and KaiB monomer [B]_tot_ are fixed to the standard condition (1.2/2 *μ*M and 3.5 *μ*M, respectively).

### States of KaiC considered in the present model

At each ATP binding site in C1, the ATP- and ADP-bound states are considered. On the other hand, states of ATP binding site in C2 are distinguished by phosphorylation of Ser431 and Thr432 and bound nucleotide. They are denoted by

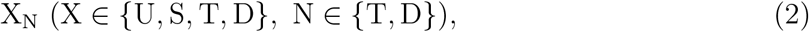

where X represents a phosphorylation state, i.e. U (unphosphorylated), S (only Ser431 is phosphorylated), T (only Thr432 is phosphorylated), or D (doubly phosphorylated), and subscript N represents a bound nucleotide, i.e. T (ATP) or D (ADP).

In the present model, one C1-specific and two C2-specific conformational transitions are considered. C1 undergoes a conformational transition between the KaiB-binding-competent (BC) and -incompetent (BI) conformational states. On the other hand, C2 undergoes conformational transitions between the rigid and flexible conformational states, and between the buried and exposed conformational states. Hence, four combinational states, i.e. rigid-buried, rigid-exposed, flexible-buried, and flexible-exposed states are considered in C2 (in the present study, the rigid-exposed state will be omitted later).

In the present model, we considered all possible combination of the states listed above. The present model consists of a set of rate equations of elementary reactions that connect the combinational states.

### Processes at each ATP binding site in C1 in the present model

At each ATP binding site in C1, we consider ATP hydrolysis and ADP/ATP exchange. We assume that the phosphate release after the ATP hydrolysis is rapid and refer to the combination of these two processes simply as ATP hydrolysis. We also assume that the incorporation of exogenous ATP after the release of bound ADP is rapid and refer to the combination of these two processes as ADP/ATP exchange.

The rate constant of the ATP hydrolysis, *k*_1h_, is assumed to be independent of any KaiABC states for simplicity. On the other hand, the ADP/ATP exchange is allowed only when C2 is in the flexible state and C1 is in the BI state. We denote the rate constant of the ADP/ATP exchange in this case by 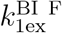 and in other case by 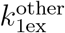, which is fixed to 0 in the present study.

### Conformational transition of C1 in the present model

We assume that the rate constants of the conformational transition from BI to BC state, *k*_1IC_, depends on the number of bound ADP in C1 *n*_1ADP_. Following the Monod-Wyman-Changeux model for multimerization effects[40] and the previous models for KaiABC oscillator[28, 29], we represent the dependence by the exponential form as

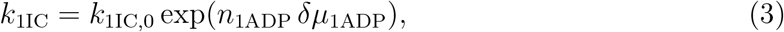

where *k*_1IC,0_ is the rate constant when C1 fully binds ATP, and *δμ*_1ADP_ is the factor for free energy difference. Moreover, *δμ*_1ADP_ is restricted to be positive to describe the promotion of the conformational transition from BI to BC state by the accumulation of ADP in C1.

On the other hand, we assume that the rate constants of the conformational transition from BC to BI state, *k*_1CI_ is independent of any KaiABC states for simplicity.

### (De)formation of C1-KaiB-KaiA complex in the present model

In the present model, at most six KaiB monomers can bind to C1 of a KaiC hexamer only when C1 is in the BC conformational state, and a KaiA dimer can bind to one of the KaiB monomers bound to C1. We assume that these bindings are much faster than other processes and adopt the rapid equilibrium approximation for these bindings. The equilibrium constants of the KaiB and KaiA bindings are denoted by *K*_1B_ and *K*_1A_, respectively. For simplicity, *K*_1B_ and *K*_1A_ are assumed to be independent of the number of KaiB bound to C1 or the number of KaiA bound to KaiB. Furthermore, the conformational transition of C1 from BC to BI state is assumed to occur regardless of the KaiAB binding, followed by the immediate release of the KaiA dimers and KaiB monomers on C1 as expressed by

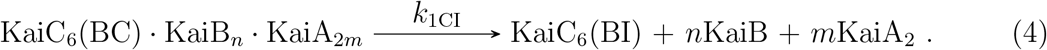

Under the above assumptions, the total concentration of KaiC hexamer in the BC conformational state binding *n* KaiB monomers, [C_6_(BC) *·* B_*n*_]_tot_, is given by

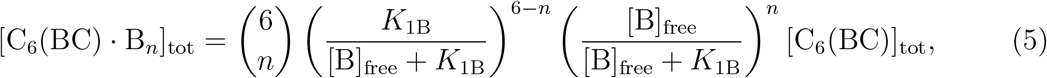

where [C_6_(BC)]_tot_ is the total concentration of KaiC hexamer in the BC conformational state, and [B]_free_ is the concentration of unbound KaiB monomer. The concentration of C_6_(BC) *·* B_*n*_ binding *m* KaiA dimers, [C_6_(BC) *·* B_*n*_ · A_2*m*_], is given by

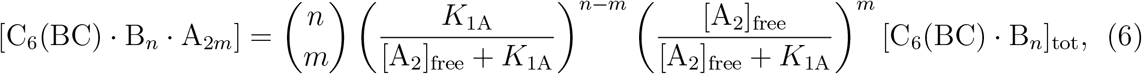

where [A_2_]_free_ is the concentration of unbound KaiA dimer. The total concentration of KaiB monomer bound to C1, [B]_onC1_, is given by

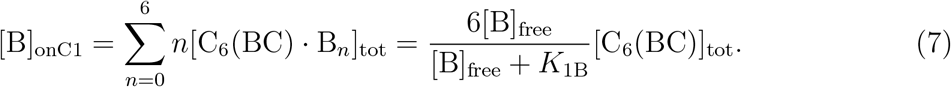

Moreover, the total concentration of KaiA dimer bound to the KaiB-C1 complex is given by

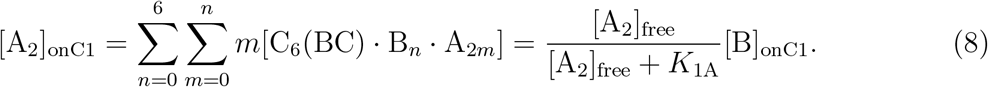

Following the previous models of the KaiABC oscillator[8, 25, 28, 29], we assume that the bindings of KaiB and KaiA are very strong. In the present study, we fix both *K*_1B_ and *K*_1A_ to 1.0 *×* 10^*−*5^*μ*M. Under the standard condition with this *K*_1B_, almost all KaiC hexamers in the BC conformational state bind six KaiB monomers.

### Processes at each ATP binding site in C2 in the present model

At each ATP binding site in C2, we consider ATP hydrolysis and ADP/ATP exchange as in C1 (Fig. 1). Moreover, we consider phosphorylation and dephosphorylation reactions of Ser431 and Thr432 (Fig. 1), in which a phosphate is transferred between the bound nucleotide and either of Ser431 and Thr432.

For simplicity, we assume that the rate constant of the ATP hydrolysis, *k*_2h_, is independent of any KaiABC states. On the other hand, we assume the ADP/ATP exchange is allowed only when C2 is in the exposed conformational state. We denote the rate constant of the ADP/ATP exchange in this case by 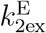 and when C2 is in the buried state by 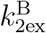, which is fixed to 0 in the present study.

We assume that the rate constants of Ser431 phosphorylation and dephosphorylation between U_T_ and S_D_ (*k*_S+_ and *k*_S*−*_, respectively) depend on the rigid/flexible conformational states of C2 as well as the rate constants of the phosphorylation and dephosphorylation of Thr432 between U_T_ and T_D_ (*k*_T+_ and *k*_T*−*_, respectively). This dependence is explicitly expressed by superscripts R (rigid) and F (flexible) as in 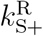. For simplicity, the rate constants of the phosphorylation and dephosphorylation between T_T_ and D_D_ are assumed to be the same as those between U_T_ and S_D_, and those between S_T_ and D_D_ are also assumed to be the same as those between U_T_ and T_D_.

The all processes in each C2 ATP binding site are summarized in Eq. (9).

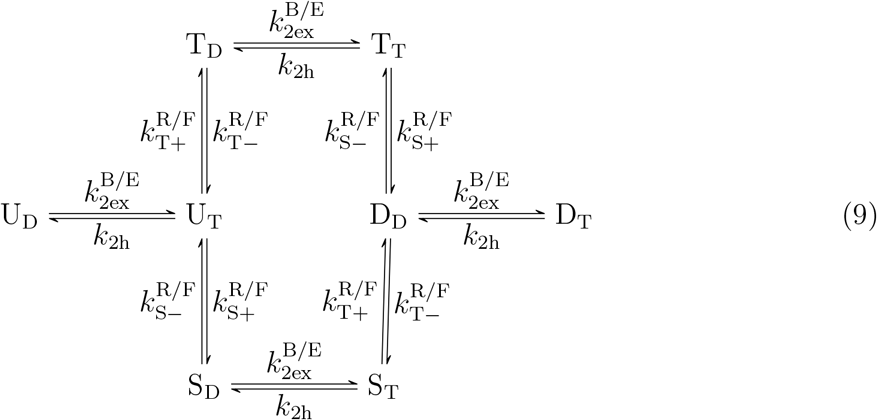

### Conformational transitions of C2 and KaiA binding to C2 in the present model

In the present model, we assume that at most one KaiA dimer can bind to C2 of a KaiC hexamer only when C2 is in the exposed conformational state. We also assume that the KaiA binding to C2 and the two conformational transitions of C2 are much faster than other processes and adopt the rapid equilibrium approximation for these processes.

The equilibrium constant between the buried and exposed conformational states, *K*_2EB_, is assumed to depend on whether C2 is in the rigid or flexible conformational state 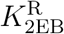 and 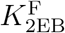, respectively). On the other hand, the equilibrium constant between the rigid and flexible conformational states, *K*_2FR_, is assumed to depend on whether C2 is in the buried or exposed conformational state (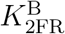 and 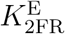, respectively). Moreover, we assume that 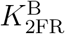 and 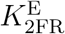 depend on the phosphorylation-nucleotide states of C2. We represent the dependence by an exponential form as

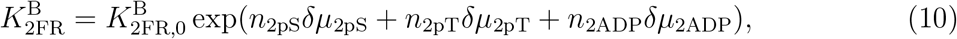

where *n*_2pS_, *n*_2pT_, and *n*_2ADP_ denote the numbers of the phosphorylated Ser431 and Thr432, and the number of bound ADP in C2, respectively; *δμ*_2pS_*, δμ*_2pT_, and *δμ*_2ADP_ are the corresponding factor for free energy difference; and 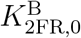 is the equilibrium constant when C2 is fully dephosphorylated and fully binds ATP. Note that the rigidification of C2 by the phosphorylation of Ser431 is described by positive *δμ*_2pS_. In the present model, 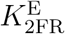 is determined by the detailed balance as explained below (Eq. (17)). For simplicity, we assume that the equilibrium constant of the KaiA binding to C2, *K*_2A_, as well as 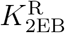 and 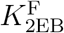 is independent of the phosphorylation-nucleotide states of C2. Note that even under this assumption, the affinity between KaiA and C2 still depends on the phosphorylation-nucleotide states because the population of the exposed state, which is competent of KaiA binding, is controlled by the phosphorylation-nucleotide dependence of the rigid/flexible conformational transition.

The conformational transitions of C2 and KaiA binding to C2 are summarized in Eq. (11).

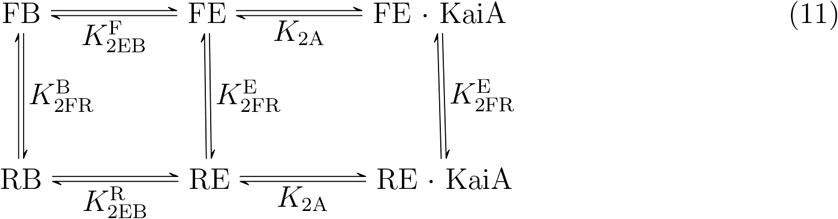

In the schematic representation described above, XY (X *∈ {*F, R*},* Y *∈ {*E, B*}*) represent the conformational state of C2, i.e. F, R, E, and B denote the flexible, rigid, exposed, and buried conformational states, respectively.

In the present study, we further assume that the exposed conformational state is absent when C2 is in the rigid state, that is

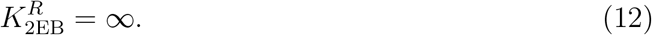

In this case, the scheme in Eq. (11) is reduced to

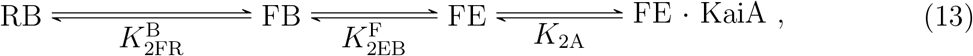

which is equivalent to the scheme shown in Fig. 2B. The total concentration of KaiA dimer bound to C2 is given by

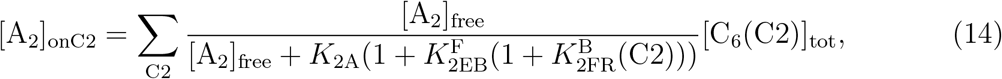

where 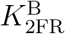 (C2) and [C_6_(C2)]_tot_ are 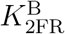 and the total concentration of KaiC hexamer in a C2’s phosphorylation-nucleotide state.

### Conservations of total KaiB monomer and KaiA dimer in the present model

The conservations of the total concentrations of KaiB monomer [B]_tot_ and KaiA dimer [A_2_]_tot_ are represented as

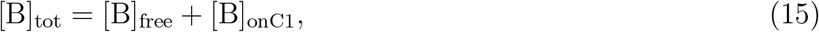

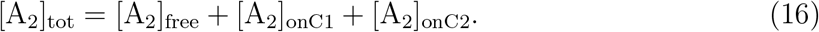

Thus, [B]_free_ and [A_2_]_free_ can be obtained by solving Eqs. (15) and (16) with Eqs. (7), (8), and (14).

### Constraints arising from the detailed balance in the present model

To assure thermodynamic consistency, the present model imposes the detailed balance condition on the processes constituting a loop. As in Paijmans’s model[29], we omit the ATP hydrolysis and the ADP/ATP exchange from the processes subjected to the detailed balance. Note that the allosteric communication between C1 and C2 in the present model is unrelated to the detailed balance because the present allostery is represented only by the rate constant of ADP/ATP exchange in C1.

For the loop consisting of the conformational transitions between the rigid and flexible conformational states and between the buried and exposed conformational states (Eq. (11)), the detailed balance is given by

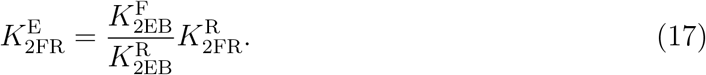

In the present model, there exists a loop consisting of the (de)phosphorylation of Ser431 and the conformational transitions between the rigid and flexible conformational states as shown in Eq. (18).

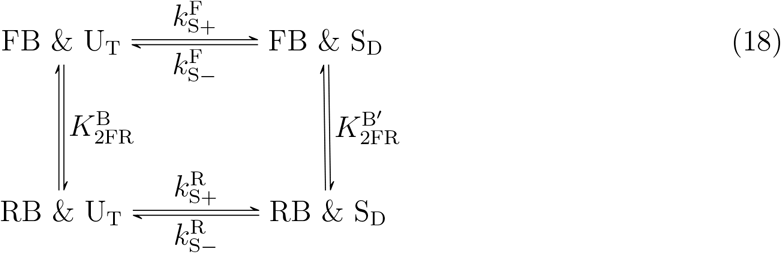

The detailed balance for this loop is

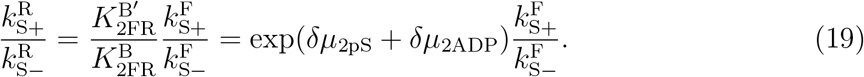

Similarly, the detailed balance about Thr432 is expressed as

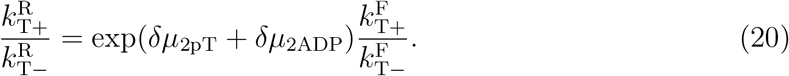

### Temperature dependence of parameters in the present model

We assume an Arrhenius temperature dependence for all the constants in the present model. The rate and equilibrium constants are represented by two parameters, the pre-exponential factor determined at 30 °C and the activation energy for a rate constant or the activation energy difference for an equilibrium constant. A rate constant *k*(*T*) and an equilibrium constant *K*(*T*) are represented as

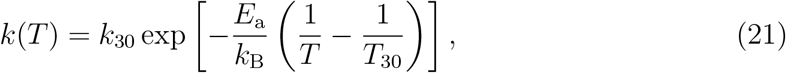

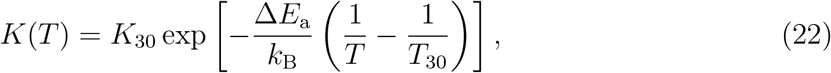

respectively, where *E*_a_ is the activation energy of the rate constant, Δ*E*_a_ is the activation energy difference of the equilibrium constant, *k*_30_ and *K*_30_ are the values at 30 °C of the corresponding constants, *T*_30_ is the temperature at 30 °C (= 303.15 K), and *k*_B_ is the Boltzmann constant. Factors for free energy difference in Eqs. (3) and (10) are represented by

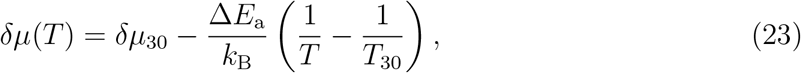

where *δμ*_30_ is the value *δμ* at 30 °C.

### Parameter Optimization

Optimization of the rate and equilibrium constants in the present model is performed to reproduce several experimental data including their temperature dependence. The parameter optimization is conducted through an automatic minimization of the difference between the experimental data and the outputs of the model. The loss function to be optimized considers multiple types of incorporates data simultaneously as

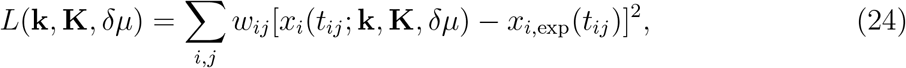

where *x_i,_*_exp_(*t_ij_*) and *x_i_*(*t_ij_*; **k**, **K***, δμ*) denote the *i*th experimental data at time *t_ij_* and the corresponding output of the model with parameters **k**, **K**, and *δμ*, respectively, and *w_ij_* is the weight.

In the present study, the term *x_i_*(*t_ij_*; **k**, **K***, δμ*) was obtained through the integration of the rate equations by the fourth-order Runge-Kutta method, and the loss function was automatically minimized by the Nelder-Mead method.

The following experimental data were used in the parameter optimization:

- Phosphorylation in the presence of KaiAB at 26-40 °C (every 4 °C) from time *t* = 0 h to *t* = 120 h (every 3 h) [37];
- Phosphorylation in the presence of KaiA at 30 °C at time *t* = 20 h [8];
- Phosphorylation (only U and T states) in the absence of KaiAB at 30 °C at time *t* = 20 h [38];
- Ratio of ATP in bound nucleotide (only maximum and minimum values) in the presence of KaiAB at 30 °C [16];
- Ratio of ATP in bound nucleotide in the presence of KaiA at 30 °C at time *t* = 8 h [16];
- Ratio of ATP in bound nucleotide in the absence of KaiAB at 30 °C at time *t* = 8 h [16];
- The maximum and minimum values of ATPase activity (i.e., rate of ADP production) in the presence of KaiAB at 30 °C [41];
- ATPase activity in the presence of KaiA at 30 °C at time *t* = *∞* [41];
- ATPase activity in the absence of KaiAB at 30 °C at time *t* = *∞* [41];
- Q10 values of ATPase activity in the absence of KaiAB at time *t* = 0.1 and *t* = *∞* [22];
- Minimum value of [B]_free_ (in the presence of KaiAB at 30 °C) restrained to 0.38 *μ*M to reproduce the arrhythmic/rhythmic phase transition around [B]_tot_ *∼* 1.75 *μ*M [3]; and
- Period when [A]_tot_ *∼* 3.6 *μ*M [3].

In the calculation of the phosphorylation oscillation at various temperatures, the initial state of KaiC was set to the steady state (*t* = *∞*) of KaiC in the absence of KaiAB at 0 °C.

The optimized parameters are shown in Table I.

## DATA AVAILABILITY

All data generated in this study are available from the corresponding authors upon request.

## ACKNOWLEDGEMENTS

The authors are grateful to Y. Fruike, J. Abe, A. Mukaiyama, and S. Akiyama for the provision of raw experimental data in Ref. [37]. This work has been supported by JSPS KAKENHI, Grant Number JP18K14185 (SK) and JP16H02254 (SS), and the Indo-Japan bilateral collaboration program. The calculations were partially carried out on computers at the Research Center for Computational Science, Okazaki, Japan.

## AUTHOR CONTRIBUTIONS

S.K. and S.S. designed research. S.K. performed research. S.K. and S.S. analyzed data and wrote the paper.

## ADDITIONAL INFORMATION

Supplementary information is available for this paper.

