## Supplemental Information for "Sufficiency of unidirectional allostery in KaiC in generating the cyanobacterial circadian rhythm"

### Supplementary Information for "Sufficiency of unidirectional allostery in KaiC in generating the cyanobacterial circadian rhythm"


Shin-ichi Koda* and Shinji Saito†

*Department of Theoretical and Computational Molecular Science, Institute for Molecular Science, 38 Nishigo-Naka, Myodaiji, Okazaki, Aichi 444-8585, Japan*



(Dated: April 2, 2020)



*

†

#### SUPPLEMENTAL TEXT

##### Results of the fitting

Figure S1 shows the results of the fitting in the parameter optimization. The phosphorylation oscillations of KaiC at 28, 32, 36, and 40 °C shown in Figs. S1A-D are in good agreement with the corresponding experimental data[S1], as well as Fig. 3A-D. In the present model, the ratio of ATP in bound nucleotide at 30 °C in the presence of KaiA and KaiB (Fig. S1E) is also in good agreement with the experimental results[S2]. Figure S1F shows the ATPase activities at 30 °C. Although the magnitude of the ATPase activities obtained from the model slightly deviate from the experimental data[S3], the present model can reproduce the asymptotic behaviors, such as the saturation of the activity in the presence of KaiA.


[S1] Y. Furuike, J. Abe, A. Mukaiyama, and S. Akiyama, Biophysics and Physicobiology **13**, 235 (2016).

[S2] T. Nishiwaki-Ohkawa, Y. Kitayama, E. Ochiai, and T. Kondo, Proceedings of the National Academy of Sciences **111**, 4455 (2014).

[S3] K. Terauchi, Y. Kitayama, T. Nishiwaki, K. Miwa, Y. Murayama, T. Oyama, and T. Kondo, Proceedings of the National Academy of Sciences **104**, 16377 (2007).

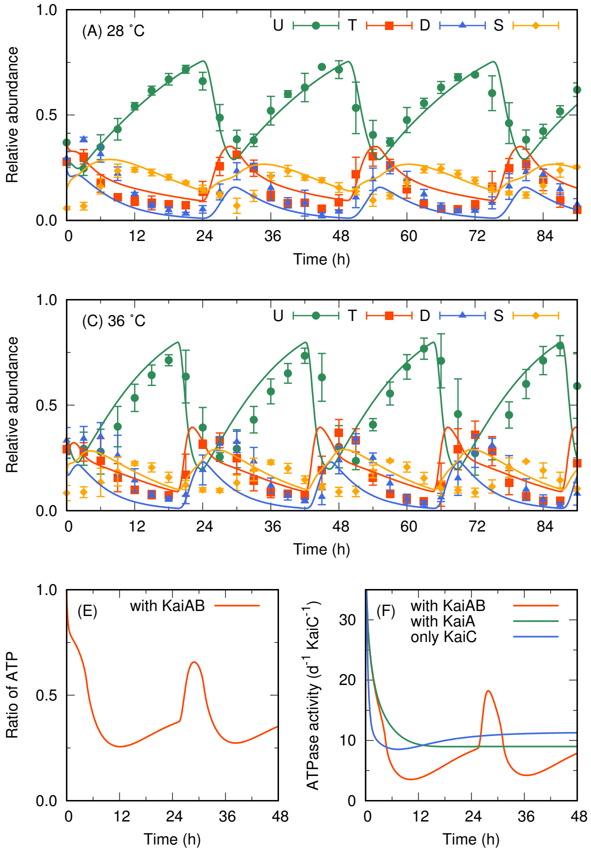


FIG. S1. Results of the present model with optimized parameters. Solid curves and dots represent the results of the model and the experimental data used in the parameter optimization. (A-D) Relative abundance of phosphorylation states (U, T, D, and S) of KaiC in the presence of KaiA and KaiB at 28, 32, 36, and 40 °C. Dots and error bars indicate the experimental data and their standard deviations[S1], respectively. The initial state of the model is the steady state in the absence of KaiAB at 0 °C. (E) Ratio of bound ATP in KaiC to bound nucleotide at 30 °C in the presence of KaiA and KaiB. The initial state is fully dephosphorylated and binds ATP. (F) ATPase activities at 30 °C. The initial state is the same as that used in (E).
